# Inexperienced bumble bees are poor at securely landing regardless of flower orientation or presence of labellum

**DOI:** 10.1101/2024.06.27.601070

**Authors:** S.R. McWaters, S. Popp, M. D. Rivera, F. Mendoza, A. Dornhaus

**Affiliations:** Ecology and Evolutionary Biology, University of Arizona; Department for Mathematics and Computer Science, Free University Berlin

**Keywords:** foraging, labellum, flower orientation, landing, pollination

## Abstract

The mutualism between bees and flowers creates strong selection on both the structure of the flower and behavior of the bee to maximize pollination and foraging success, respectively. Previous research has primarily assessed the costs of foraging by quantifying the time and accuracy of search, and handling time of the flower. However, there is little attention given to the actual success of landing, and it is often not explicitly stated whether failed landing attempts are taken into consideration. We show here that landing attempts often are unsuccessful, especially in inexperienced bees. Orientation of artificial flowers in our experiment neither influenced the preference nor landing success of a naive bumble bee forager. The presence of a labellum, often considered to serve as a landing platform, also did not influence landing success, indicating that it may mostly play a role in flower recognition or act as a nectar guide. Failed landing attempts may thus play an under-recognized role in the foraging efficiency and behavior of bees, and learning may be key in individual bee landing efficiency, not just flower recognition. Further research should aim to quantify the costs of landing failures and consider the role of experience in individual bee landing success.

---

Plants and their respective pollinators have been subject to intense coevolutionary pressures that have resulted in many adaptations of plants and pollinators improving pollination for the plant and energy gain for the pollinators (1,2). Bees are a primary pollinating group for many flowering plants due to the generalist nature of many species (3,4). Any aspect of foraging that influences foraging success has likely been evolutionarily selected for especially in social bees, as even a small benefit per flower visit will translate to a significant benefit over a bee’s lifetime and for the colony as a whole (5).

Most research on floral structure and pollinator foraging either focuses on the process of locating flowers, i.e. the visual and cognitive aspects of flower choice before landing, or on flower handling, i.e. the time from successfully landing on the flower to the bee accessing the reward (rev in Chittka and Spaethe, 2007). Between flower choice and flower handling, bees must first land on the flower. Landing is one of the more under-studied aspects of bee foraging behavior, with some recent studies focusing on flight control, including in the context of landing (7–9). However, there seems to be a gap in our knowledge of what role landing plays in overall foraging, and whether the process of landing itself is a challenging task: i.e., do bees fail at it, and is such failure costly? What is the role of learning in achieving a smooth landing? We know that flower morphology influences bee choice from a visual stance, but does it also affect the landing success? While flower orientation has been shown to influence preferences (10,11) there is little to no information on how landing changes depending on the direction the flower faces. It seems intuitive to think that the orientation of the flower would affect landing success. For example, labellums, which are enlarged petals that stick out horizontally amongst vertically oriented flowers, are well-known nectar guides and are commonly referred to as landing platforms even though we lack empirical evidence that they aid in pollinators successfully landing on a flower.

Here, we aim to understand how the orientation and morphology (the labellum) of a flower aids in pollinator foraging, namely in preference and landing success in naive bumble bees (*Bombus impatiens*). Bumble bees are ideal models as they are generalist pollinators and naturally visit a wide variety of flowers with varying orientations and morphologies (12). They are easily maintained in a lab and readily forage from artificial flowers. Using artificial flowers, we can directly and independently manipulate morphological floral features, while keeping visual signals similar. Artificial flowers are commonly used in pollinator foraging experiments (13–16). We manipulated either the orientation of the flower face (experiment 1) or the presence of a labellum (experiment 2). We test whether orientation of the flower influences the bee’s ability to successfully land; eg., bees might be more successful landing on horizontally oriented flowers (flat, facing up). We also test whether bees show a preference for one of these flower morphologies. If structures such as labellums act as a landing platform for pollinators, we expect that bees will preferentially land on labellums rather than other flower parts, and be more successful in such landings than on the vertically oriented petals or flowers without a labellum.

## Methods

### Bee maintenance

Small (10-50 worker) *Bombus impatiens* bumble bee colonies were purchased from Koppert Biological Systems, Michigan. They were maintained in wooden nest boxes (39 × 10.5 × 23cm) connected via a plastic tube to a larger flight arena (76.2 × 59.7 × 39.4cm). On non-experimental days the colonies were fed sugar solution from a multi-well feeder raised off the ground in the flight arena. All workers in the colonies were uniquely marked with a numbered color tag fixed to the thorax with super glue. Newly emerged workers were marked daily.

### Defining ‘Landing success’

In both experiments, we measured individual foragers’ preferences and success in landing on arrays of artificial flowers. We define a landing attempt as a bee touching the flower with one or more legs. A landing attempt was scored as successful when the bee remained securely on the flower without falling. This meant coming to a complete stop and ceasing to beat the wings while attached to the flower with all legs. Successful landings were generally followed by the forager crawling around the flower to gain access to the nectar reward. Examples of unsuccessful landings are the bee brushing past a petal, or resuming flying or falling off after the first contact.

### Assays

Thirty minutes prior to testing, the colony was fed from the artificial flowers (Experiment 1) or multi-well feeder (Experiment 2) in the flight arena to stimulate foraging activity. The identity of foragers on the feeder(s) was noted. After 30 minutes, all individuals were manually removed from the flight arena and the access tube blocked. Identity of foragers that approached the tube was noted for later testing, but only one individual was allowed into the flight arena at a time. A bee that entered the flight arena was then given 10 minutes in which to make her first landing attempts before she was removed from the flight arena. If the individual did not fly at all, it was removed after 5 minutes. Individuals that made landing attempts were allowed to naturally return to the hive (via attempts to return through the tube) or were removed after 10 minutes passed from their last landing attempt on a flower. Individuals who had made landing attempts were removed from the colony, however those that made no attempts were allowed to return into the hive.

### Experiment 1: Flower Orientation

These simple artificial flowers consisted of yellow craft foam disks (5 cm in diameter) with a microcentrifuge-tube-well in the center, to hold a reward of 5 µl (Fig. 1a). The flowers were elevated on a wire stem (∼15 cm) stuck in a black rubber stopper as base. Wire stems of some flowers were bent to give the surface of the ‘flower’ a vertical orientation. Tested individuals were exposed to 10 vertical and 10 horizontal flowers simultaneously. Each quadrant of the flower array contained either 2 horizontal and 3 vertical or 3 horizontal and 2 vertical flowers. Within each quadrant these were placed randomly. The array of flowers was replaced and washed after each individual was tested.

**Figure 1.**
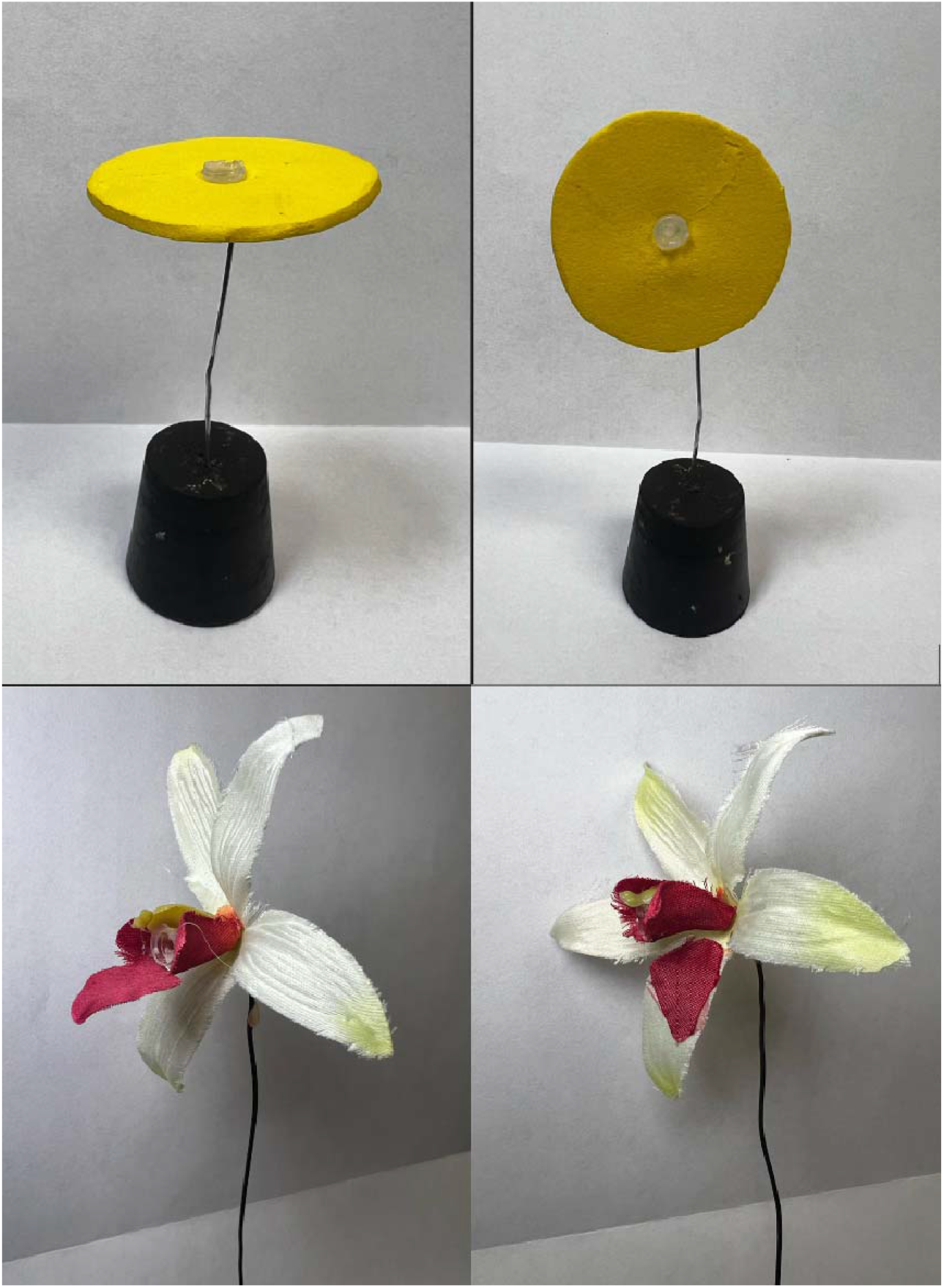
Artificial flowers used. (a) Vertical (top left) and horizontal (top right) flowers used in experiment 1. (b) Flowers with the labellum attached (bottom left) and detached (bottom right), as used in experiment 2.

### Experiment 2: Presence of a labellum

We used silk flowers from a local craft shop (Michael’s Stores Inc.) roughly resembling a simple orchid flower to construct artificial flowers for this experiment (Fig. 1b). The flowers had white outer petals and red, partially fused petals forming a tube in the middle. This tube provided a place to hold a microcentrifuge tube with the nectar reward. The tube ended in a large bright red labellum, which was either left intact (Flowers with labellum) or removed and attached to an outer (white) petal (Flowers without labellum). Thus, the two flower types were similar in size and color pattern to an approaching bee, but one provided a possible ‘landing platform’ and the other did not.

The flowers were elevated in the same manner as in Experiment 1 by attaching them to a wire stem, while ensuring that all landing platforms were at the same height and angle. Foragers were presented with an array of four flowers, two of each type in alternating order. The array was placed in the flight arena opposite to the nest access. To stimulate foraging activity on the flowers, we also applied ∼1µl of linalool oil on an inaccessible spot underneath the microcentrifuge tube for training and test trials.

## Results

### Experiment 1: Flower orientation

A total of 57 bees were observed. Over all landing attempts (n = 169), regardless of flower type, bees were successful 73 percent of the time. Overall, we saw no pattern of preference for horizontal or vertical flowers (Wilcoxon-Test, p = 0.64, n = 84 landing attempts on horizontal and 85 on vertical flowers). This remains true when looking at only the first landing attempt for each bee (Wilcoxon-Test, p = 0.15, n = 23 on horizontal and 34 on vertical flowers). The orientation of the flower (vertical or horizontal) did not influence average success of landing at each flower type (mean success rates: vertical = 0.70, horizontal = 0.73; Wilcoxon-Test, p = 0.21; Figure 2A). Again the same was true when looking only at first landing attempts for each bee (mean success rates: vertical = 0.64, horizontal = 0.78; Chi-square, p = 0.31).

**Figure 2.**
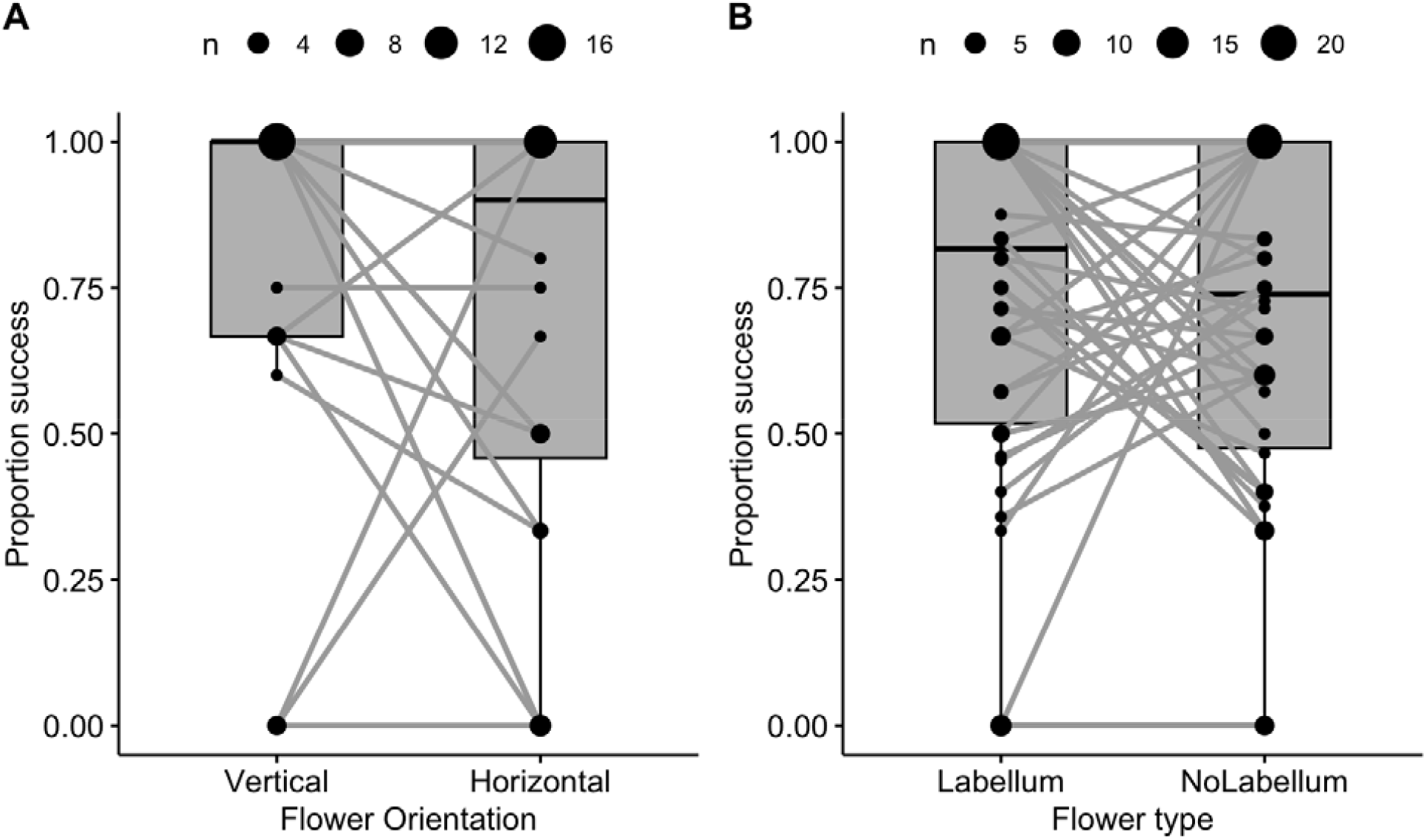
**(A)** Proportion of landings that were successful for each bee, and median proportion across all bees, on either vertical or horizontal flowers. We saw no significant difference in landing success between flower types (n = 57 bees, Wilcoxon-test = 0.21). Dark gray lines between box plots represent individual bees and the size of the points indicates how many individuals are represented. **(B)** Proportion of landing attempts that were successful for each bee on flowers with and without a labellum. We saw no significant difference in landing success between flower types (n = 80, Wilcoxon-test p = 0.38). Lines between box plots represent individual bees, point size indicates the number of individuals represented..

### Experiment 2: Presence of labellum

A total of 66 bees were observed. Over all landing attempts (n = 454), regardless of flower type, bees were successful 66 percent of the time. Overall, we saw no pattern of preference for flowers with a labellum or without (Wilcoxon-Test, p = 0.42, n = 221 landing attempts on labellum flowers and 223 on non-labellum flowers). This observation remains true when looking at only first landing attempts (Wilcoxon-Test, p = 0.81, n = 32 on labellum flowers and 34 on non-labellum flowers). The presence of a labellum did not influence success of landings overall (mean success rate: labellum = 0.73, no labellum = 0.69; Wilcoxon-Test, p = 0.38; Figure 2B), nor among first landing attempts for each bee (mean success: labellum = 0.78, no labellum = 0.76; chi-square, p = 1). A linear model of success rate and thorax width showed that bees with larger thorax widths were overall less successful at landings (n = 29 bees, R^2^ = 0.06, t = -2.214, p = 0.03; Figure 3). To determine if the presence of a labellum played a role in the success of these larger bees we split each bee’s success rate between flower types and ran a linear model to test the interaction between thorax width and flower type on landing success. The model shows that this interaction was not significant in the landing success of bees (n=29 bees, R^2^ = 0.01, t = 0.410, p = 0.68; Figure 3).

**Figure 3.**
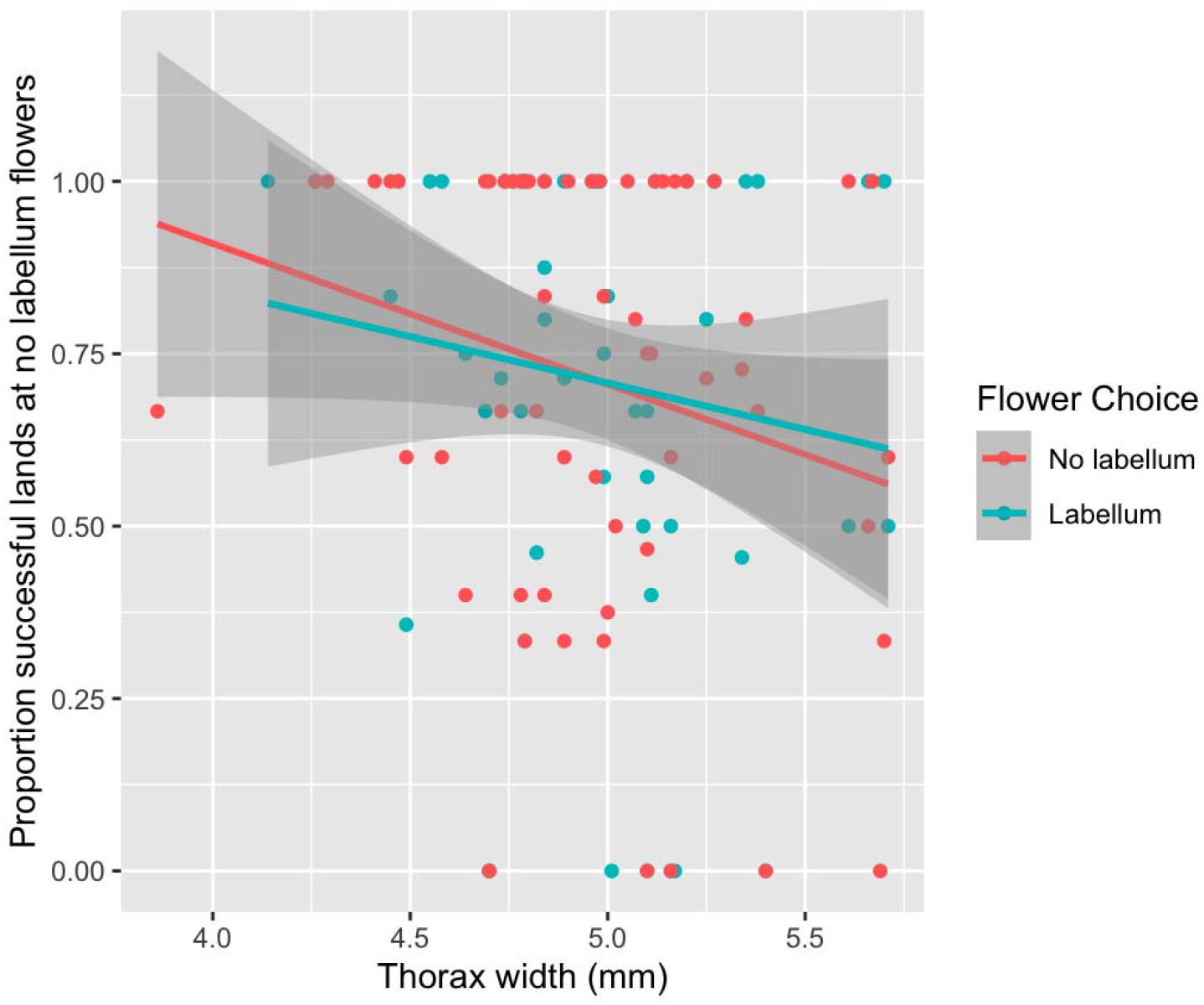
Proportion of successful lands (y-axis) and thorax width (x-axis) for each individual bee at flowers without a labellum (red; n = 59) and with a labellum (blue; n = 56). Shaded regions are standard error. Thorax width was a significant factor in predicting landing success (linear model: n = 29 bees, R^2^ = 0.06, p = 0.03) but the type of flower did not play a role (linear model: n = 29 bees, R^2^ = 0.01, p = 0.68).

## Discussion

Here, we observed that bumble bees do often fail their landing attempts and this is especially true for bees with larger thorax widths. Most literature is focused on the location and handling of flowers (17,18) and some on the mechanics of flight during the approach to landing (8,19,20), but the landing attempt itself and its consequences on foraging needs further attention. Our experiment shows that inexperienced bees often fail at landing. Bees are likely to improve with experience, which was not measured in this study. Bees in the wild are likely to face many more environmental factors that may make landing unpredictable or difficult (e.g. wind, light, temperature). For example, light intensity was shown to influence the timing of leg extension when approaching a flower in bumble bees but not in sweat bees but the success of the landing attempt was not specifically measured (19).

*Bombus impatiens* bumble bees in our study did not prefer either vertically or horizontally oriented flower corollas, nor flowers with or without a ‘landing platform’ (labellum). In addition to this lack of preference, we found no evidence that the orientation of the flower or presence of a labellum aids in landing success, despite the fact that bees often did not land successfully. The indifference in preference between horizontal and vertical flowers stands in contrast to previous studies using real flowers that found bees were more likely to land on vertically facing flowers (11,21). Our conflicting findings in comparison to the literature may potentially be attributed to the fact that the symmetry of flowers used in our experiment did not differ. In nature, there is a correlation in flower morphology between orientation and symmetry (22). Flowers which possess only one (vertical) bilateral symmetry axis (‘zygomorphic’) are mostly vertically oriented (22) and bumble bees have shown a preference for bilateral over radial symmetry (23). It is thus important to test how flower symmetry and orientation interact to shape bee preferences, in order to get a more comprehensive understanding of flower preference in pollinators.

The functionality of labellums has been largely linked to the mutualism between flowers and pollinators. Research has shown that the labellum patterns are sometimes sexually deceptive, tricking male pollinators into visitation by resembling a female (24). Additionally, it is extremely common in the literature to refer to the labellum as a ‘landing platform’ (24–27) without having empirical evidence supporting this claim. Our findings suggest we may need to re-think our assumption that the labellum increases the ease of landing but further research should test this on a variety of pollinators.

## Acknowledgements

We would like to thank Sofia Tsai and Nicole Richards as well as the STAR laboratories for their contribution to data collection. FM was funded by the University of Arizona GPSC grant. We also thank the NSF (DBI 1564521 to AD) and the EEB Department for funding.

